# Dynamic Histone Lysine Methylation and Demethylation in Wood Frog (*Rana sylvatica*) Liver During Anoxia

**DOI:** 10.64898/2026.07.05.736536

**Authors:** Pallabi Chakraborty, Kenneth B. Storey

## Abstract

Anoxia is a major stress for most vertebrates and frequently accompanies harsh winter conditions, particularly in species that spend much of the season frozen solid. North American freeze-tolerant wood frogs (*Rana sylvatica*) can survive several months without oxygen and endure whole-body freezing for up to eight months of the year, with ∼70% of total body water frozen as extracellular ice, yet revive when temperatures rise in spring. Survival depends on multiple adaptations, including tolerance of prolonged oxygen deprivation while frozen, when breathing and circulation are halted. A key strategy involves hepatic glycogen mobilization, producing large amounts of glucose that are distributed to tissues where it functions both as a cryoprotectant and as a substrate for anaerobic ATP production.

The present study examines the role of histone lysine methylation and demethylation in regulating liver proteins under anoxic conditions. Relative protein expression of seven histone methyltransferases (ASH2L-S, ASH2L-L, RBBP5, SETD8, SMYD2, ESET, SETD1), six lysine demethylases (KDM1A, KDM3B, KDM4A, KDM4B, KDM5A, KDM5C), and eight histone marks (H3K4me1, H3K4me2, H3K9me3, H3K27me3, H3K36me3, H3K79me3, H4K20me1, H4K20me3) were evaluated in wood frog liver under control, 4-hour, and 24-hour anoxia exposures. The data indicate that histone lysine methylation and demethylation contribute significantly to transcriptional regulation under anoxia. Specifically, H3K4, H3K36, and H3K79 methylation were associated with transcriptional activation, whereas H3K9, H3K27, and H4K20 methylation correlated with transcriptional repression. These findings highlight the dynamic role of epigenetic regulation in supporting hypometabolism and stress adaptation in freeze-tolerant wood frogs.

## Introduction

All cells, tissues, and organs of vertebrate bodies typically contain the same genes, with a few exceptions such as red blood cells and platelets. However, gene expression levels differ widely among cells, tissues, and organs, often under the influence of epigenetic controls. Epigenetic modifications begin during embryonic development and are central to cell, tissue, and organ diversification, while also mediating responses to both internal metabolic needs and external stresses. Importantly, epigenetic regulation allows environmental factors and behavior to alter gene expression without changing DNA sequences, responding readily to influences such as stress, pollution, chemicals, diet, aging, and drugs. According to the National Institutes of Health (USA, 2009) in their Epigenomics Initiative: “Epigenetics refers to both heritable changes in gene activity and expression (in the progeny of cells or individuals) and stable, long-term alterations in the transcriptional potential of a cell that are not necessarily heritable.” In this study, we focus on histone lysine methylation and demethylation as key epigenetic mechanisms that contribute to transcriptional regulation and stress adaptation in the freeze-tolerant wood frog (*Rana sylvatica*) under anoxia.

One of the most remarkable animal adaptations to environmental stress is freeze tolerance—the ability of certain vertebrate and invertebrate species to endure long-term whole-body freezing, with as much as 65–70% of total body water frozen in extracellular and extra-organ spaces (Storey & Storey, 2017a). This phenomenon has been documented in a variety of terrestrial invertebrates as well as several amphibian and reptile species inhabiting high latitudes or altitudes. Among amphibians, the North American wood frog (*Rana sylvatica*) is the best-studied model for exploring survival strategies. These frogs overwinter on the forest floor beneath leaf litter and snowpack, which provides insulation from the lowest winter temperatures, yet they must still endure months in a frozen state. Survival requires strict metabolic regulation to withstand prolonged whole-body freezing without breathing or blood circulation, relying primarily on anaerobic glycolysis to meet ATP demands.

During freezing, wood frogs face multiple life-threatening stresses, including the suspension of vital physiological functions such as breathing, heartbeat, muscle contraction, and blood circulation, as well as the formation of extracellular and extra-organ ice crystals that can compress or puncture cells. As a result, metabolism, digestion, waste filtration, and other essential processes are halted (Kling et al., 1994; Layne & First, 1991; Storey & Storey, 2004a). Long-term exposure to subzero temperatures further disrupts metabolic balance. Excessive ice formation in extracellular spaces stops heartbeat and breathing, blood volume declines due to water loss into ice masses, and the cessation of circulation quickly leads to tissue hypoxia and anoxia. Remarkably, wood frogs can withstand chronic hypoxia for several months even without freezing (Holden & Storey, 1997), a condition that would be fatal to humans within minutes (Watanabe & Morita, 1998). Freeze-tolerant species also possess mechanisms to minimize or prevent damage from ice formation. In non-tolerant species, ice growth can rupture capillary walls or damage cells that shrink extensively due to water loss or mechanical stress from expanding extracellular ice crystals (Rubinsky et al., 1987). By contrast, wood frogs employ protective strategies that reduce such injuries, enabling survival under prolonged freezing conditions.

Wood frogs employ a range of molecular adaptations to survive winter in a frozen state. These include entry into a hypometabolic state, activation of cellular stress responses to anoxia and dehydration (i.e., water loss into extracellular ice masses), and pre-hibernation accumulation of glycogen reserves that provide glucose for anaerobic catabolism. Anoxia tolerance is further supported by maximizing anaerobic ATP production, minimizing acidosis, and upregulating anoxia-specific genes, proteins, and enzymes (Storey & Storey, 2017b). Anoxia is one of the major stress factors that wood frogs must endure to survive in the frozen state. To cope, they employ two primary adaptations: (1) production of high concentrations of organic osmolytes, primarily glucose, which provide both a large supply of fuel for anaerobic metabolism and colligative protection to minimize cell volume reduction and prevent intracellular water from freezing, and (2) restriction of ice formation to extracellular and extra-organ compartments within the body (Storey & Storey, 2017). Freezing halts respiration and blood flow, leading to tissue anoxia, but wood frogs counter this by mobilizing large amounts of glucose that function simultaneously as cryoprotectant and substrate for anaerobic glycolysis—the main ATP-generating pathway available to cells in frozen animals (Gupta & Storey, 2021).

Suppression of energy-expensive pathways, such as protein synthesis, is a key feature of anoxia tolerance in wood frogs. In addition, these animals activate anoxia-specific genes and produce stress-responsive proteins, including ice-nucleating proteins and glycolipids. Enzyme activity is further reduced through post-translational modifications, while epigenetic mechanisms such as DNA methylation, histone modifications, and miRNA-mediated inhibition of mRNA transcription provide additional regulatory control. Cytoprotective systems—including molecular chaperones, antioxidant defenses, and anti-apoptotic pathways—are also upregulated as part of the winter survival strategy (Storey & Storey, 2017). A central component of hypometabolism is the suppression of highly energy-demanding processes such as transcription and translation, while maintaining selective upregulation of crucial genes (Bocharova et al., 1992; Fraser et al., 2001; Storey & Storey, 2017a; Frerichs et al., 1998; Van Breukelen & Martin, 2001). These processes are among the costliest in cells—for example, the formation of a single peptide bond requires approximately 5 ATP (Storey & Storey, 2004b)—and are therefore downregulated under stress or hypometabolic conditions (Storey & Storey, 1990). Nevertheless, activation of selected genes remains essential to synthesize protective proteins that safeguard wood frogs during freezing, anoxia, or dehydration. These include stress-responsive proteins that promote cellular stability, along with the downregulation of genes involved in oxygen-dependent processes and oxygen-sensitive signaling pathways (Storey & Storey, 2017d; Wu et al., 2018).

Metabolic rate depression (MRD) is a critical survival strategy for many animal species, particularly during winter hibernation or diapause (Storey & Storey, 2004c). Wood frogs employ MRD to endure sub-zero temperatures by reducing flux through energy-expensive biochemical pathways. The liver plays a central role in this process: as freezing begins, its large glycogen reserves are rapidly mobilized, producing high concentrations of glucose that are exported via the bloodstream to other organs. This glucose functions both as a cryoprotectant, protecting cells from freezing damage, and as a substrate for anaerobic metabolism. Wood frogs tolerate ice formation in extracellular and extra-organ spaces, while ice-nucleating proteins help regulate the rate and extent of ice crystal growth, ensuring controlled freezing that minimizes cellular injury.

Epigenetic mechanisms can significantly alter gene expression patterns through cytosine methylation, post-translational modifications of histone proteins and chromatin remodeling, and RNA-based regulation (Gibney & Nolan, 2010). When blood plasma freezes, most metabolic activity is suspended due to the lack of oxygen and nutrient delivery to tissues and organs. Ice crystals form in extracellular spaces, puncturing blood vessels and causing internal bleeding, while the withdrawal of water from cells leads to dehydration and shrinkage. Remarkably, wood frogs can withstand the loss of up to 60% of total body water into extracellular ice during freezing (Churchill & Storey, 1993). Blocking water supply to cells not only disrupts volume maintenance and structural integrity but also halts nutrient transfer and waste removal. As freezing begins, ice rapidly forms in the abdominal cavity and around internal organs, while the liver mobilizes large glycogen reserves to produce glucose. This glucose is exported into the bloodstream for rapid uptake by other organs, serving both as a cryoprotectant and as fuel for anaerobic metabolism in the frozen state. By allowing ice to form outside cells and organs while preventing intracellular freezing, wood frogs avoid the lethal damage suffered by most organisms exposed to subzero temperatures.

DNA methylation and histone modifications play crucial roles in survival under stress conditions. In particular, histone lysine methylation and demethylation are closely interconnected with DNA methylation and contribute to epigenetic regulation of key biological processes, including cell cycle control, development, differentiation, and responses to DNA damage. Methyl groups can be attached to specific amino acids on the N-terminal “tails” of histone proteins, influencing how tightly DNA is wound around the histone core. This structural modulation of chromatin directly impacts transcription, either promoting gene activation or enforcing repression. The nucleosome core is composed of four histones—H2A, H2B, H3, and H4—with two molecules of each forming the histone octamer, which is further stabilized by H1/H5 linker histones. These core histone proteins contain a globular carboxyl (C) domain and a flexible amino (N) domain that facilitates the addition of post-translational modifications (PTMs). The N-terminal tails of H3 and H4 in the nucleosome octamer play a critical role in chromatin remodeling, often mediated by lysine methyl-binding proteins (Hyun et al., 2017). Transcriptional modification of histone tails is closely linked to chromatin organization, thereby regulating gene expression. Chromatin exists in two major states: euchromatin, which is less condensed, gene-rich, and transcriptionally active, and heterochromatin, which is highly condensed, gene-poor, and transcriptionally silent, as demonstrated by ChIP-seq analyses.

Histone lysine methylation is mediated by histone lysine methyltransferase (KMT) enzymes, which primarily target lysine residues on core histones H3 and H4 to regulate DNA accessibility and thereby activate or suppress gene transcription (Rea et al., 2000). Most methyltransferases contain a conserved Su(var)3-9, Enhancer of Zeste, Trithorax (SET) domain that catalyzes methyl group transfer using S-adenosyl-L-methionine (SAM) as the donor. Regulation of transcription is further coordinated by “writer,” “reader,” and “eraser” proteins: writers (KMTs) add methyl marks, erasers (lysine demethylases, KDMs) remove them, and readers recognize and bind the modified residues to mediate downstream effects (Black et al., 2012a; Zhang & Reinberg, 2001). The functional outcome of lysine methylation depends on the degree of modification, with mono-, di-, or tri-methylation (me1, me2, me3) exerting distinct effects that can either promote transcriptional activation or enforce repression (DeLange et al., 1973).

The present study focuses on the action of lysine methyltransferases (KMTs) that activate transcription by modulating H3K4, H3K36, and H3K79 residues, as well as the associated lysine demethylases (KDMs) that repress transcription by removing methyl marks at H3K9, H3K27, and H4K20 (Black et al., 2012b). H3K4 methylation is controlled by methyltransferases in the SET1/MLL1 (SET1A and SETD7) and MLL2/KMT2B complexes, which are the main writers of H3K4 methyl marks associated with gene activation. For example, SET1A binds with retinoblastoma protein 5 (RBBP5) and ASH2-like histone methyltransferase (ASH2L) (Li et al., 2023a; Qu et al., 2018). Trimethylation of H3K4 (H3K4me3) marks transcriptional start sites of active genes, promoting transcription through recruitment of PHD-domain proteins. By contrast, H3K4me1 and H3K4me2 are linked with enhancer elements and transcriptionally active genes. H3K36me3 acts as a transcriptional activator regulated by SET2 and SMYD2. The SET domain is responsible for catalysis, adding or removing methyl groups (CH3) from SAM. Similarly, H3K79 di- or tri-methylation (me2/3) by the MLL complex is essential for maintaining enhancer–promoter interactions (Godfrey et al., 2021). KMTs form dynamic complexes that regulate chromatin and nucleosome arrangement in many organisms (Li et al., 2023b).

By contrast, repressive histone modifications such as H3K9 methylation exhibit diverse interactions with the nucleosome and are regulated by the KMT1 family, including SUV39H1 (Padeken et al., 2022). H3K9 is trimethylated by SUV39H1 and ESET, maintaining transcriptional silencing, with heterochromatin protein 1 (HP1) acting as a reader of H3K9me3 to reinforce repression (Lomberk et al., 2006). HP1 also influences euchromatin organization and interacts with other post-translational modifications. Similarly, H3K27me3 is associated with repression of cell type-specific genes, while H4K20 methylation—catalyzed by SETD8, the sole enzyme in the KMT5 family—contributes to suppressive effects at the nucleosome level (Martin & Zhang, 2005; Nishioka et al., 2002). In the wood frog brain, repressive H3K9me and H3K27me marks were reduced during recovery from freezing, correlating with changes in several KMTs (Bloskie & Storey, 2022). These findings highlight the dynamic regulation of histone methylation as a key mechanism in stress adaptation.

The present study investigates the relative protein levels of seven histone lysine methyltransferases (KMTs)—SMYD2, SET8, RBBP5, ASH2L-S, ASH2L-L, ESET, and SETD1A—and six lysine demethylases (KDMs: KDM1A, KDM3B, KDM4A, KDM4B, KDM5A, and KDM5C) to assess their responses to 4-hour and 24-hour anoxia exposures. In addition, changes in eight histone marks were evaluated: H3K4me2, H3K4me3, H3K9me3, H3K27me3, H3K36me3, H3K79me3, H4K20me1, and H4K20me3. The study focuses on the wood frog liver, a tissue highly susceptible to stress and known to undergo adaptive changes in histone lysine methylation and demethylation, which are considered key regulators of gene expression under anoxia. Epigenetic regulation plays a critical role in stress tolerance, yet its contribution to hypometabolism during anoxia remains poorly understood. Previous studies have emphasized transcriptional controls as central to metabolic suppression, but the specific epigenetic mechanisms in wood frogs have not been fully defined. The objective of this study was therefore to investigate histone modifications and related epigenetic mechanisms in the liver of *Rana sylvatica* under anoxia stress. We hypothesize that epigenetic factors associated with transcriptional activation are downregulated, while those linked to transcriptional repression are upregulated under anoxia, thereby supporting hypometabolism.

This study is the first to investigate active histone lysine methylation and demethylation in the liver of Rana sylvatica under anoxia, addressing a gap left by previous work that focused mainly on DNA methylation, microRNA regulation, and proteomic responses.

## Materials And Methods

### Animal collection

Adult male wood frogs (*Rana sylvatica*) weighing 5-7 g were captured from breeding ponds near Ottawa, Ontario, Canada during the early spring season. In the lab, frogs were given a tetracycline bath and then housed in plastic boxes containing damp sphagnum moss. After a period of acclimation at 5°C for two weeks (without feeding), control frogs were selected from this group. Other frogs were exposed to 4 h or 24 h anoxia treatment at 5°C, following a procedure previously described (Gerber et al., 2016). Briefly, the animals were transferred to plastic jars lined with damp paper towels (pre-wetted with distilled water that had previously been bubbled with 100% nitrogen gas) and held in crushed ice. Jars were then flushed with nitrogen gas for 20 min via ports in the lids. Frogs were then quickly added to the jars (4-5 per container), and jars were again flushed with nitrogen gas for 20 min followed by closing input and output ports and sealing the lids with parafilm. Jars were returned to 5°C for 4 h or 24 h. After anoxia exposure, jars were reconnected to the nitrogen gas lines and frogs were quickly sampled after 4 h or 24 h anoxia exposures. All frogs were euthanized by pithing and liver was immediately dissected from control, 4 h anoxia and 24 h anoxia condition, flash frozen in liquid nitrogen, and stored at -80°C until use. All animal experimental procedures had the prior approval of the Carleton University Animal Care Committee (protocol no. 106935) and followed the guidelines of the Canadian Council on Animal Care.

### Total protein isolation

Total protein was isolated from samples of frozen liver (∼100 mg each) from 5 different individuals for each condition (control, 4h anoxia, 24h anoxia) as previously described (Al-attar et al., 2019; Hawkins & Storey, 2018). After weighing, tissues were crushed with a mortar and pestle under liquid nitrogen. Frozen samples were then quickly homogenized 1:5 w:v with a P10 homogenizer in ice-cold lysis buffer (EMD Millipore 43-045) and combined with 1 mM Na_3_VO_4_, 10 mM NaF, 10 mM of B-glycerophosphate, and 10 µl/ml of protease inhibitor cocktail containing 104 mM AEBSF, 80 µM aprotinin, 4 mM bestatin, 1.4 mM E-64, 2 mM leupeptin, 1.5 mM pepstatin A (Bioshop, P1C001.1).

Tissue homogenates were centrifuged at 10,000 rcf for 15 min at 4°C and then the supernatant containing soluble proteins was collected. Protein concentration of each sample was determined using the BioRad protein assay (Catalogue #500-0002; BioRad Laboratories, Hercules, CA, USA) at 595 nm on an MR5000 microplate reader (Dynatech Laboratories, Chantilly, VA, USA). Subsequently, all samples were standardized to 10 μg/μL protein by adding calculated small volumes of homogenization buffer. Samples were then mixed 1:1 v: v with 100 mM-Tris buffer pH 6.8 containing sodium dodecyl sulfate (4% w/v SDS), 20% v/v glycerol, 0.2% w/v bromophenol blue, and 10% v/v 2-mercaptoethanol final concentrations of 5 µg/µl protein per sample. All samples were then boiled in a water bath for 10 min to fully denature and linearize the proteins. Samples were then stored at −40 °C until use.

### Nuclear protein isolation

Cytoplasmic and nuclear extracts were performed following a modified version of the protocol described by Al-attar and Storey (2020). Nuclear protein samples were prepared from frozen liver tissue samples (weighing ∼50-100 mg) from control, 24 h frozen and 8 h thawed frogs (n = 4 biological replicates). Tissue samples were homogenized with a Dounce homogenizer 1:5 w/v in ice cold 1X cytosolic fraction buffer containing 10 mM HEPES buffer pH 7.9, 10 mM KCl, 10 mM EDTA, 20 mM β-glycerophosphate, 10 μL/ml of 100 mM dithiothreitol (DTT) and 10 μL/mL of protease inhibitor cocktail (Bioshop, Cat # PIC001.1) followed by brief vortexing and then incubation on ice for 60 min. Homogenates were then centrifuged at 12,000 rpm for 15 min at 4°C and supernatants were collected (cytoplasmic fraction). The pellets were washed in cytosolic buffer and centrifuged once again under the same conditions (12,000 rpm for 15 min at 4°C). Pellets were then collected and re-suspended in 5X nuclear fraction buffer containing 100 mM HEPES, 2 M NaCl, 5 mM EDTA, 50% (v/v) glycerol, 100 mM β-glycerophosphate, with 10 μl of 100 mM DTT and 10 μl protease inhibitor cocktail, pH 7.9, followed by sonication and then incubation on ice for 10 min. Samples were then centrifuged at 14,000 rpm for 10 min at 4°C and supernatants were collected (nuclear fraction). Protein concentrations were determined using the Bradford assay (Bio-Rad, Cat # 500-0002). Samples were tested via immunoblotting with a cytoplasm-specific marker, α-tubulin (Abclonal, Cat # AC007) and a nucleus-specific marker, histone H3 (Cell Signaling Cat # 9715) to validate the separation of cytoplasmic and nuclear fractions.

### Western immunoblotting

Protein samples of equal amounts from liver of control, 4 h anoxia, and 24 h anoxia frogs (15–40 μg/ml, depending on target protein to be detected) were loaded into 6–15% SDS-polyacrylamide gels (the % acrylamide in the resolving gel depended on the molecular weight of the protein being probed). Molecular weight markers were also loaded in other lanes, 3 μL of BLUeye prestained protein ladder (10-245 kDa; Catalogue #PM007-0500; FroggaBio, Toronto, ON, Canada).

The upper stacking gel (pH 6.8) was comprised of 5% acrylamide v/v in 1 M Tris buffer with 0.1% SDS, 0.1% APS (ammonium persulphate), and 0.1% TEMED (N,N,N′,N′-tetramethylethane-1,2-diamine) added, whereas resolving gels (pH 8.8) were 8–15% acrylamide v/v in 1.5 M Tris buffer with 0.1% SDS, 0.1% APS, and 0.1% TEMED added. Loaded gels were run in a BioRad Mini Protean III system (BioRad Laboratories, Hercules, CA, USA) for 30–180 min at 180 V in running buffer (25 mM Tris-base, 190 mM glycine, 0.1% w/v SDS, pH 7.6). Proteins from the gels were then transferred onto 0.45 μm pore PVDF membranes by electroblotting at room temperature for 45-180 min at 160 mA in 1X transfer buffer (25 mM Tris-base, 192 mM glycine, 10% v/v methanol, pH 8.5). To prevent non-specific binding of primary and secondary antibodies, membranes were incubated in 1X TBST (20 mM Tris-base, 140 mM NaCl, 0.05% Tween-20) with either (a) skimmed milk, 1–10%, on a rocker at room temperature for 30 min) or (b) polyvinyl alcohol (PVA) MW 30,000–70,000 (1 mg/mL) on a rocker at room temperature for 30–90 s) in 1X TBST. Membranes were then probed with primary antibodies (1:1000 v/v dilution in 1X TBST) at 4°C overnight. Relative protein levels of nine lysine methyltransferases (KMTs), and their methylated histone marks were assessed via western blotting. Following incubation with primary antibodies, blots were washed for 3 × 5 min with 1X TBST and probed with HRP-conjugated anti-rabbit/mouse secondary antibodies (1:8000 v/v dilution in TBST; Catalogue #APA002P, BioShop Canada Inc., Burlington, ON, Canada) at room temperature for 30 min. Membranes were washed again for 3 x 5 minutes in 1X TBST and were then visualized by chemiluminescence (1:1 v/v H2O2 and Luminol) using a ChemiGenius Bio Imaging System (Syngene, Frederick, MD, USA). Membranes were then stained with Coomassie blue (0.25% w/v Coomassie brilliant blue, 7.5% v/v acetic acid, 50% methanol) to visualize all protein bands for loading standardization.

### Statistical analysis

Chemiluminescent protein bands on immunoblots were quantified by densitometry using the ChemiGenius Bio Imaging System and GeneTools Software (Syngene, Frederick, MD, USA). Band densities were standardized against the combined intensity of Coomassie blue stained bands in the same lane that did not show differential expression between conditions and that were well separated from the immunoblot band of interest. Data for each experimental condition are expressed as mean ± SEM for n = 4 samples from different animals. Statistical analysis was performed using a one-way ANOVA and Tukey’s post-hoc test with p<0.05 accepted as a significant difference and using the RBioPlot statistical package (J. Zhang & Storey, 2016).

## Results

### ASH2L and SETD8 are anoxia-responsive lysine methyltransferases

Relative protein expression levels of seven KMTs (ASH2L-S, ASH2L-L, RBBP5, SETD8, SMYD2, ESET, and SETD1A) were measured in liver of *R. sylvatica* comparing protein levels from control versus with anoxia exposure (4 h and 24 h) (Fig. 1) via Western blotting. RBBP5 and SMYD2 did not show significant changes under the three conditions. By contrast, ASH2L-S and ASH2L-L showed significant changes under anoxia. After 24h anoxia, ASH2L-S levels were significantly increased to 1.39 ± 0.09 whereas ASH2L-L levels were decreased to 1.22 ± 0.04. While in 4h anoxia, ASH2L-L level returned to above control levels to 1.42 ± 0.06 but ASH2L-S has fallen by 30% (1.09 ± 0.06) than 24h but remained up from control. SETD8 showed a different pattern with a decrease of 30% i.e. (0.71 ± 0.14-fold) under 4 h anoxia as compared with controls but after 24h, anoxia level was significantly increased by 50% (1.51 ± 0.18-fold) over controls, respectively, and were almost doubled compared with the 4 h anoxia values. ESET and SETD1A remained same in all three conditions.

**Figure 1.**
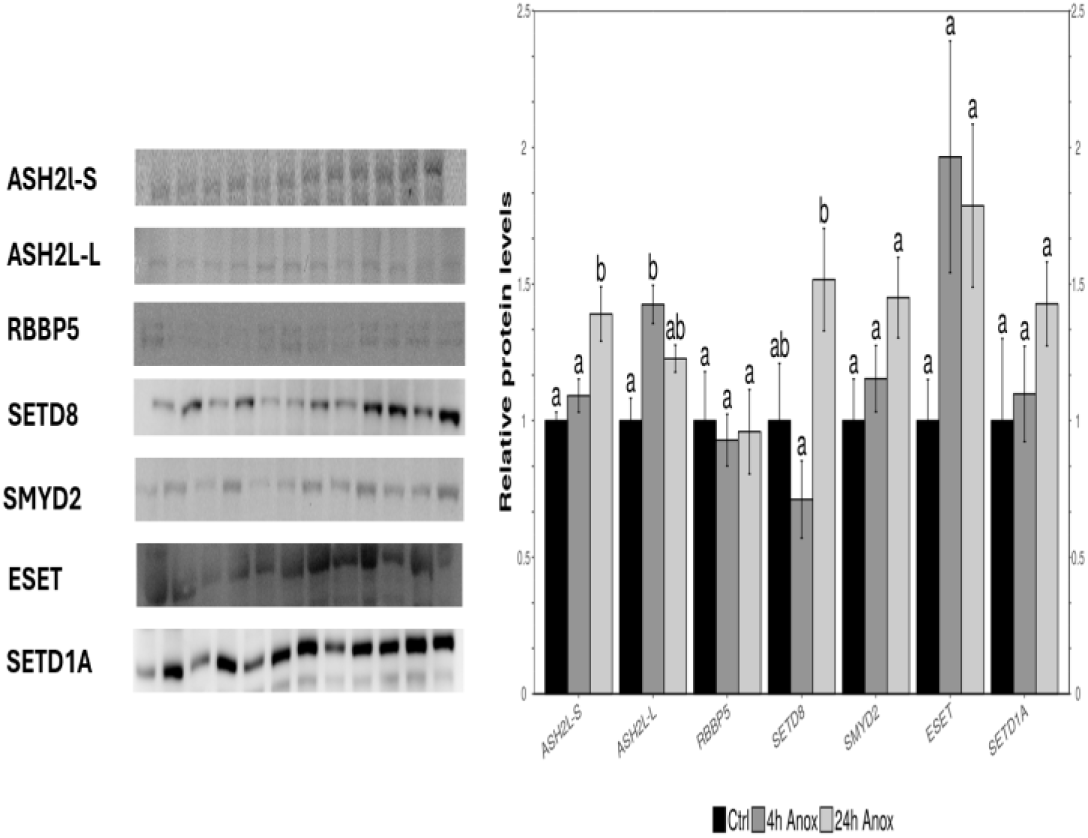
Effects of 4 h or 24 h anoxia exposure on the relative levels of 7 proteins (ASH2L-S 60kDa, ASH2L-L 80kDa, RBBP5, SETD8, SMYD2, ESET, and SETD1A methyltransferases) in extracts of *R. sylvatica* liver. Histogram shows the relative levels of histone lysine methylation pathway regulators including the methyltransferases (SET8 and SMYD2) and associated complex proteins (RBBP5, ASH2L-S, and ASH2L-L) in extracts of wood frog liver comparing responses to 4 h and 24 h anoxia with aerobic controls as detected by Western blotting. Data mean SEM± SEM, n=4 for control, 4 h or 24 h anoxia samples. Data were assessed using analysis of variance with p<0.05 accepted as a significant difference. Bars labeled “a” or “b” indicate significant differences.

### Analysis of methylated histone residue levels

Relative protein levels of 8 histone lysine methyl marks (H3K4me2, H3K4me3, H3K9me3, H3K27me3, H3K36me3, H3K79me3, H4K20me1, and H4K20me3) in liver of wood frog (Fig. 2) were measured under control, 4 h and 24 h anoxic conditions. H3K27me3 and H4K20me3 were the only marks that changed significantly under anoxia, decreasing to just 0.18 ± 0.07 and 0.14 ± 0.06 of control values during 4 h anoxia exposure. However, in both cases, values for both histone marks rose again after 24 h anoxia exposure. H3K27me3content rose back to a value of 0.86 ± 0.42 as compared to controls whereas H4K20me3 level increased by 1.33 ± 0.25fold but remained 20% lower as compared to control. Both H4K20me3 and H3K27me3 marks are strongly related to silencing of gene expression. Responses by H3K4me2, H3K4me3, H3K9me3, H3K36me3, H3K79me3 and H4K20me1 also showed the same pattern, decreasing during 4h anoxia and increasing again after 24h anoxia although these changes were generally not different from either control or 24 h anoxia conditions.

**Figure 2.**
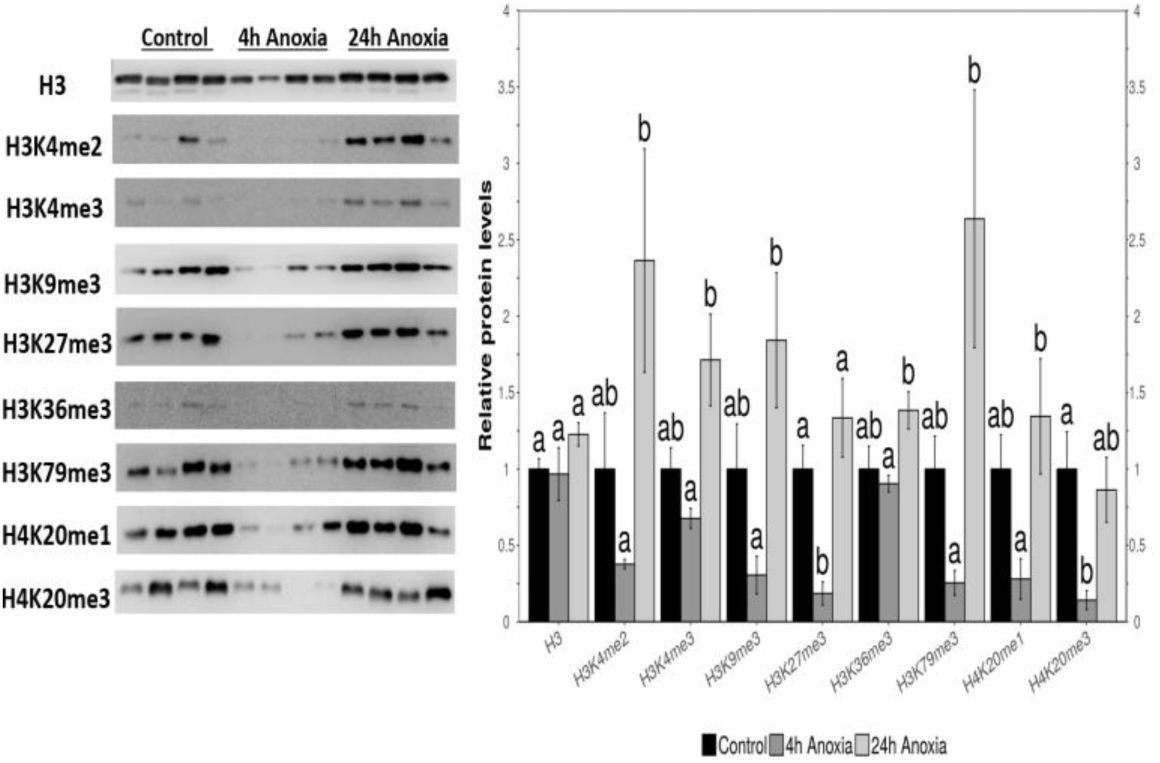
Effects of 4 h anoxia and 24 h anoxia on the relative protein levels of H3 total protein and eight histone modifications: H3K4me2, H3K4me3, H3K9me3, H3K27me3, H3K36me3, H3K79me3, H4K20me1 and H4K20me3 in extracts of wood frog (*R. sylvatica*) liver as detected by Western blotting. Data means ± SEM, n=4 (control, 4 h anoxia and 24 h anoxia). Data were analyzed using analysis of variance (p<0.05 accepted as a significant difference). Bars labeled “a” or “b” indicate significant differences.

### KDM protein expression in response to anoxia

The responses to 4 h and 24 h anoxia exposure by lysine demethylases (KDMs) were also assessed (Fig. 3). Compared with controls, protein expression was assessed in response to 4 h and 24 h anoxia conditions in wood frog liver. Relative protein expression of six histone lysine demethyltransferase proteins (KDM1A, KDM3B, KDM4A, KDM4B, KDM5A, KDM5C) were also measured in wood frog liver comparing controls with 4 h and 24 h anoxia. There were no significant changes observed between these three conditions in wood frog liver under anoxic conditions. However, lysine demethylation is possible, when lysine demethylases (KDMs) get all four conditions such as oxygen, iron, oxoglutarate, and ascorbic acid. Under anoxic conditions (absence of oxygen) error bars look same but may be some molecular activity could be possible.

**Figure 3.**
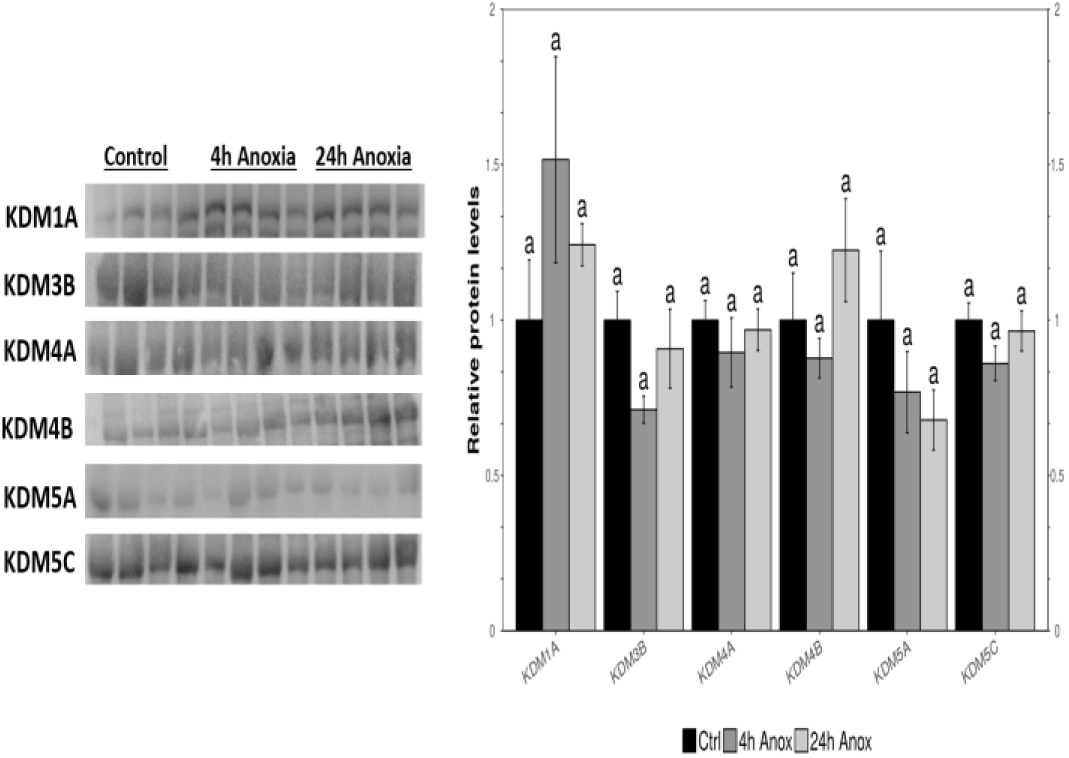
Effects of 4 h anoxia and 24 h anoxia on the relative protein levels of KDM1A, KDM3B, KDM4A, KDM4B, KDM5A, KDM5C lysine demethylation in extracts of wood frog (R. sylvatica) liver as detected by Western blotting. Data means ± SEM, n=4 for control, 4 h anoxia and 24 h anoxia conditions. Data were analyzed using analysis of variance (p<0.05 accepted as a significant difference). Bars “a” labeled or “b” indicate significant differences.

## Discussion

The main cause of wood frogs to survive in harsh winter by depressing their metabolic rate so they reduce their expensive energetic demand to support their limited carbohydrate stores during hibernation especially when there is a scarcity of food and moreover body is totally frozen. Therefore, wood frogs suspend important functions such as heartbeat, breathing, muscle movement etc. that in turn cells experience anoxia during frozen state, thus anoxic tolerant species need a very well-organized metabolic process to support the anoxia (Storey and Storey, 2017). Furthermore, several cryoprotective and energy saving pathways are also involved to keep down the damage during anoxic condition (Lung and Storey, 2022). A balance between ATP production and ATP utilization is essential for the survival of wood frog. Approximately, 32 ATP produce through oxidative phosphorylation in the presence of oxygen in mitochondria whereas in absence of oxygen i.e. anaerobiosis only 2 ATP produce, again anaerobiosis is an energetically costly because it provides energy 5 to 10 times more than aerobic levels but the process deplete energy quickly. Also, production of lactic acid cause animal fatigued (Laurie Vitt et al., 2009).

Therefore, histone modifications are key epigenetic regulatory mechanisms that have important roles in many cellular functions that support anoxic environment. However, histone lysine methyl transferases (KMTs), and demethylases play a crucial role in gene transcription or suppression as well as chromatin modulation. To understand how histone methylation is regulated under anoxic conditions, we focused our study on histone lysine methylation and demethylation in *R. sylvatica* liver so that wood frogs can use ATP in a well-organized way. SETD8 is related to a number of cellular activities that involve cell cycle regulation, DNA replication, DNA damage repair, and gene expression via H4K20 methylation (Beck et al., 2012, Jorgensen et al., 2013). H4K20 mediated by SETD8 requires genomic integrity. Recently discovered information states that SETD8 and its corresponding H4K20 methylation repair double strand breaks (DSBs) pathway (Xu et al., 2022) which may help anoxic wood frogs during early torpor and arousal state. On the other hand, ASH2L methylate H3K4 by interacting with RBBP5 and DPY30 (MLL complex) is an important activation of transcription.

The term fight or flight can be used to describe the adaptive measures used by different species to survive under adverse environmental conditions. Wood frogs chose to fight by utilizing metabolic rate depression (MRD) and other biochemical adaptations (e.g. synthesis of cryoprotectants) in order to survive months in a frozen state. Simple pathways for ATP production (e.g. glycolysis) are fueled by high reserves of glycogen in all organs to support cell/tissue survival in the frozen state. The present study also implicates various posttranslational modifications of proteins to modulate cellular metabolism in the anoxic state. One of these is histone lysine methylation/demethylation regulated by the actions of histone lysine methyl transferases (KMTs) or lysine demethylases (KDMs) that attach or remove methyl groups on lysine residues of histones (as well as other proteins) in order to modulate and/or coordinate their functions during prolonged winter freezing that causes anoxia. The role of KMTs and KDMs are crucial in regulating target proteins in an epigenetic way by expressing or repressing gene expression and subsequent protein expression to aid survival of wood frogs.

In the human body for example, KMTs are known to affect various positive functions including cell division, differentiation, and tissue/organ development but are also involved with various diseases (e.g. cancer, autoimmune diseases, etc.).

Additionally, hypoxia-induced factors (HIFs) work on low oxygen and are an important oxygen-sensitive regulator of gene expression to freezing, anoxia, and dehydration (Storey JM et al., 2022). It was observed that there is a relationship between HIFs and lysine demethylases, they altered their activity and expression to extend the hypoxic signaling response. Loss of KDM4A in hypoxic conditions leads to a decrease in HIF-1a transcriptional response. KDM4A regulates HIF-1a levels through H3K9me3 and that acts as a repressor, it gathered on HIF-1A (Grzegorz Dobrynin et al., 2017). In 24h anoxic condition H3K9me3 is upregulated double as compared to the control, which may be necessary to support the anoxic condition to save ATP.

Earlier studies have documented DNA methylation, microRNA regulation, and proteomic shifts in wood frog liver, as well as histone methylation in brain tissue. In contrast, our findings uniquely demonstrate dynamic histone lysine methylation and demethylation in liver under anoxia, identifying both activating and repressive marks. This establishes histone modifications as a novel epigenetic mechanism supporting hepatic adaptation to oxygen deprivation.

## Conclusion

To support long term survival under anoxic and ischemic conditions due to whole body freezing, wood frogs have developed multicellular adaptive responses that can regulate activation or repression of many metabolic functions and support metabolic changes so that frogs can survive over long harsh winters with many vital activities halted until they thaw in the spring (Storey and Storey, 2017).

Overall, the current study sheds light on previously unexplored regulatory mechanisms that contribute to freeze survival. By highlighting the role of KMTs in gene transcription, suppression, and chromatin modulation, our work connects histone modifications to broader biological phenomena such as disease, cell development, and differentiation. This new perspective on epigenetic regulation not only advances understanding of whole-body freeze tolerance but also suggests potential avenues in cryomedicine, including strategies to enhance long-term cold storage of human tissues and organs for transplantation.

## Acknowledgments

The author gratefully acknowledges the late Professor Kenneth B. Storey for his supervision, guidance, and scientific contributions to this work.

This preprint is made available under a Creative Commons CC BY license.

## Antibody information and suppliers for Western blotting

| **Antibody** | **Company** | **Catalogue #** |
| --- | --- | --- |
| **ASH2L-S** | Cell Signaling Technology | 5019T |
| **ASH2L-L** | Cell Signaling Technology | 5020T |
| **RBBP5** |  |  |
| **SETD8** | Cell Signaling Technology | 2996T |
| **SMYD2** | Abclonal | A6474 |
| **ESET** | Cell Signaling Technology | 2196T |
| **SETD1A** | Abclonal | A1823 |
| **KDM1A** | Abgent | A1218 |
| **KDM3B** |  |  |
| **KDM4A (JMJD2A)** | Cell Signaling Technology | 5328T |
| **KDM4B (JMJD2B)** | Cell Signaling Technology | 8639T |
| **KDM5A** | Cell Signaling Technology |  |
| **H3** | Abclonal | A2348 |
| **H3K4me2** | Abclonal | A2356 |
| **H3K4me3** | Abclonal | A2357 |
| **H3K9me3** | Abcam | Ab8898 |
| **H3K27me3** | Abclonal | A2363 |
| **H3K36me3** | Abclonal | A2366 |
| **H3K79me3** | Abclonal | A2369 |
| **H4K20me1** | Abclonal | A2370 |
| **H4K20me3** | Abclonal | A2370 |

## References

Al-attar, R., Wijenayake, S., & Storey, K. B. (2019). Metabolic reorganization in winter: Regulation of pyruvate dehydrogenase (PDH) during long-term freezing and anoxia. Cryobiology, 86, 10–18. 10.1016/J.CRYOBIOL.2019.01.006

Black, J. C., Van Rechem, C., & Whetstine, J. R. (2012a). Histone lysine methylation dynamics: establishment, regulation, and biological impact. Molecular Cell, 48(4), 491–507. 10.1016/J.MOLCEL.2012.11.006

Black, J. C., Van Rechem, C., & Whetstine, J. R. (2012b). Histone lysine methylation dynamics: establishment, regulation, and biological impact. Molecular Cell, 48(4), 491–507. 10.1016/J.MOLCEL.2012.11.006

Bloskie, T., & Storey, K. B. (2022). Epigenetics of the frozen brain: roles for lysine methylation in hypometabolism. FEBS Letters, 596(16), 2007–2020. 10.1002/1873-3468.14440

Bocharova, L. S., Gordon, R. Y., & Arkhipov, V. I. (1992). Uridine uptake and RNA synthesis in the brain of torpid and awakened ground squirrels. Comparative Biochemistry and Physiology. Part B, 101(1–2), 189–192. 10.1016/0305-0491(92)90177-S

Churchill, T.A. and Storey, K.B., 1993 Metabolic responses to dehydration by liver of the wood frog - Search Results - PubMed. (n.d.). Retrieved February 6, 2024, from https://pubmed.ncbi.nlm.nih.gov/?term=Churchill+T+A+and+Storey+K+B%2C+1993+Metabolic+responses+to+dehydration+by+liver+of+the+wood+frog

Fraser, K. P. P., Houlihan, D. F., Lutz, P. L., Leone-Kabler, S., Manuel, L., & Brechin, J. G. (2001). Complete suppression of protein synthesis during anoxia with no post-anoxia protein synthesis debt in the red-eared slider turtle Trachemys scripta elegans. The Journal of Experimental Biology, 204(Pt 24), 4353–4360. 10.1242/JEB.204.24.4353

Frerichs, K. U., Smith, C. B., Brenner, M., Degracia, D. J., Krause, G. S., Marrone, L., Dever, T. E., & Hallenbeck, J. M. (1998). Suppression of protein synthesis in brain during hibernation involves inhibition of protein initiation and elongation. Proceedings of the National Academy of Sciences of the United States of America, 95(24), 14511–14516. 10.1073/PNAS.95.24.14511

Gerber, V. E. M., Wijenayake, S. and Storey, K. B. (2016). Anti-apoptotic response during anoxia and recovery in a freeze-tolerant wood frog (*Rana sylvatica*). PeerJ 4, e1834.

Godfrey, L., Crump, N. T., O’Byrne, S., Lau, I. J., Rice, S., Harman, J. R., Jackson, T., Elliott, N., Buck, G., Connor, C., Thorne, R., Knapp, D. J. H. F., Heidenreich, O., Vyas, P., Menendez, P., Inglott, S., Ancliff, P., Geng, H., Roberts, I., … Milne, T. A. (2021). H3K79me2/3 controls enhancer-promoter interactions and activation of the pan-cancer stem cell marker PROM1/CD133 in MLL-AF4 leukemia cells. Leukemia, 35(1), 90–106. 10.1038/S41375-020-0808-Y

Gupta, A., & Storey, K. B. (2021). Coordinated expression of Jumonji and AHCY under OCT transcription factor control to regulate gene methylation in wood frogs during anoxia. Gene, 788. 10.1016/J.GENE.2021.145671

Hawkins, L. J., & Storey, K. B. (2018). Histone methylation in the freeze-tolerant wood frog (Rana sylvatica). Journal of Comparative Physiology. B, Biochemical, Systemic, and Environmental Physiology, 188(1), 113–125. 10.1007/S00360-017-1112-7

Holden, C. P., & Storey, K. B. (1997). Second messenger and cAMP-dependent protein kinase responses to dehydration and anoxia stresses in frogs. Journal of Comparative Physiology. B, Biochemical, Systemic, and Environmental Physiology, 167(4), 305–312. 10.1007/S003600050078

Hyun, K., Jeon, J., Park, K., & Kim, J. (2017). Writing, erasing and reading histone lysine methylations. Experimental & Molecular Medicine, 49(4). 10.1038/EMM.2017.11

Kling, K. B., Costanzo, J. P., & Lee, R. E. (1994). Post-freeze recovery of peripheral nerve function in the freeze-tolerant wood frog, Rana sylvatica. Journal of Comparative Physiology. B, Biochemical, Systemic, and Environmental Physiology, 164(4), 316–320. 10.1007/BF00346449

Layne, J. R., & First, M. C. (1991). Resumption of physiological functions in the wood frog (Rana sylvatica) after freezing. The American Journal of Physiology, *261*(1 Pt 2). 10.1152/AJPREGU.1991.261.1.R134

Li, Y., Ge, K., Li, T., Cai, R., & Chen, Y. (2023). The engagement of histone lysine methyltransferases with nucleosomes: structural basis, regulatory mechanisms, and therapeutic implications. Protein & Cell, 14(3), 165–179. 10.1093/PROCEL/PWAC032

Lomberk, G., Wallrath, L. L., & Urrutia, R. (2006). The Heterochromatin Protein 1 family. Genome Biology, 7(7). 10.1186/GB-2006-7-7-228

Martin, C., & Zhang, Y. (2005). The diverse functions of histone lysine methylation. Nature Reviews. Molecular Cell Biology, 6(11), 838–849. 10.1038/NRM1761

Nishioka, K., Rice, J. C., Sarma, K., Erdjument-Bromage, H., Werner, J., Wang, Y., Chuikov, S., Valenzuela, P., Tempst, P., Steward, R., Lis, J. T., Allis, C. D., & Reinberg, D. (2002). PR-Set7 is a nucleosome-specific methyltransferase that modifies lysine 20 of histone H4 and is associated with silent chromatin. Molecular Cell, 9(6), 1201–1213. 10.1016/S1097-2765(02)00548-8

Padeken, J., Methot, S. P., & Gasser, S. M. (2022). Establishment of H3K9-methylated heterochromatin and its functions in tissue differentiation and maintenance. Nature Reviews. Molecular Cell Biology, 23(9), 623–640. 10.1038/S41580-022-00483-W

Rea, S., Eisenhaber, F., O’Carroll, D., Strahl, B. D., Sun, Z. W., Schmid, M., Opravil, S., Mechtier, K., Ponting, C. P., Allis, C. D., & Jenuwein, T. (2000). Regulation of chromatin structure by site-specific histone H3 methyltransferases. Nature, 406(6796), 593–599. 10.1038/35020506

South, P. F., & Briggs, S. D. (2011) Understanding the structure and function of ASH2L: http://AtlasGeneticsOncology.org/Deep/ASH2LFunctionID20097.html DOI: 10.4267/2042/46058

Storey, K. B., & Storey, J. M. (1990). Metabolic rate depression and biochemical adaptation in anaerobiosis, hibernation and estivation. The Quarterly Review of Biology, 65(2), 145–174. 10.1086/416717

Storey, K. B., & Storey, J. M. (2004a). Metabolic rate depression in animals: transcriptional and translational controls. Biological Reviews of the Cambridge Philosophical Society, 79(1), 207–233. 10.1017/S1464793103006195

Storey, K. B., & Storey, J. M. (2017a). Molecular Physiology of Freeze Tolerance in Vertebrates. Physiological Reviews, 97(2), 623–665. 10.1152/PHYSREV.00016.2016

Van Breukelen, F., & Martin, S. L. (2001). Translational initiation is uncoupled from elongation at 18 degrees C during mammalian hibernation. American Journal of Physiology. Regulatory, Integrative and Comparative Physiology, 281(5). 10.1152/AJPREGU.2001.281.5.R1374

Watanabe, T., & Morita, M. (1998). Asphyxia due to oxygen deficiency by gaseous substances. Forensic Science International, 96(1), 47–59. 10.1016/S0379-0738(98)00112-1

Zhang, J., & Storey, K. B. (2016). RBioplot: an easy-to-use R pipeline for automated statistical analysis and data visualization in molecular biology and biochemistry. PeerJ, 4(9). 10.7717/PEERJ.2436

Zhang, Y., & Reinberg, D. (2001). Transcription regulation by histone methylation: interplay between different covalent modifications of the core histone tails. Genes & Development, 15(18), 2343–2360. 10.1101/GAD.927301

